# Teeth outside the mouth: the evolution and development of shark denticles

**DOI:** 10.1101/2022.07.13.499989

**Authors:** Rory L. Cooper, Ella F. Nicklin, Liam J. Rasch, Gareth J. Fraser

## Abstract

Vertebrate skin appendages are incredibly diverse. This diversity, which includes structures such as scales, feathers, and hair, likely evolved from a shared anatomical placode, suggesting broad conservation of the early development of these organs. Some of the earliest known skin appendages are dentine and enamel-rich tooth-like structures, collectively known as odontodes. These appendages evolved over 450 million years ago. Elasmobranchs (sharks, skates, and rays) have retained these ancient skin appendages in the form of both dermal denticles (scales) and oral teeth. Despite our knowledge of denticle function in adult sharks, our understanding of their development and morphogenesis is less advanced. Even though denticles in sharks appear structurally similar to oral teeth, there has been limited data directly comparing the molecular development of these distinct elements. Here, we chart the development of denticles in the embryonic small-spotted catshark (*Scyliorhinus canicula*) and characterise the expression of conserved genes known to mediate dental development. We find that shark denticle emergence shares a vast gene expression signature with developing teeth. However, denticles have restricted regenerative potential, as they lack a *sox2+* stem cell niche associated with the maintenance of a dental lamina, an essential requirement for continuous tooth replacement. We compare developing denticles to other skin appendages, including both sensory skin appendages and avian feathers. This reveals that denticles are not only tooth-like in structure, but that they also share an ancient developmental gene set that is likely common to all epidermal appendages.

## Introduction

Vertebrate skin appendages are an incredibly diverse group of organs that adorn the integument, including scales, spines, hair, feathers, and teeth. Despite dramatic variety in both their form and function, the early development of vertebrate skin appendages is widely characterised by the emergence of an anatomical placode (Di-Poï and Milinkovitch, 2016, Cooper et al., 2017, Harris et al., 2008); a local epidermal thickening associated with conserved gene expression patterns in both the epidermis and underlying dermis. This placode constitutes the common foundation of phylogenetically distinct skin appendages (Cooper et al., 2017). Furthermore, reaction-diffusion-like dynamics are broadly considered to control the spatial distribution of placode emergence (Kondo, 2002, Sick et al., 2006, Cooper et al., 2018).

Odontodes constitute one of the earliest known vertebrate skin appendage types (Sansom et al., 1996). These hard, mineralised, tooth-like structures have evolved over the past 450 million years to include oral teeth, branchial/pharyngeal denticles (Fraser et al., 2010), and modified enamel/dentine derived scales in certain clades of bony fishes (Mori and Nakamura, 2022, Chen et al., 2020). Furthermore, odontodes include hard, mineralised scales (or ‘skin-teeth’) that adorn the bodies of elasmobranchs (sharks, skates, and rays), known as dermal denticles. These denticles consist of a hard outer layer of hyper-mineralised enamel-like tissue (Gillis and Donoghue, 2007), a dentine layer, and a central pulp cavity, making them structurally homologous to vertebrate teeth (Fraser et al., 2010). The denticles of adult sharks facilitate numerous functions, including hydrodynamic drag reduction during locomotion, the provision of defensive armour, and communication via the binding of luminescent photophores (Oeffner and Lauder, 2012, Wen et al., 2015, Reif, 1985). Consequently, the dermal denticles of elasmobranchs have evolved to exhibit various shapes and sizes, both within and across species (Motta et al., 2012, Gabler-Smith et al., 2021). Patterns of morphological variation in shark denticles are also observable across deep time (Sibert and Rubin, 2021). Although the early development and patterning of shark denticles has been previously characterised (Cooper et al., 2017, Cooper et al., 2018), the molecular basis of their development, morphogenesis, and final morphological diversity is not yet comprehensively understood.

The elasmobranch dentition is renowned for its prolific conveyor belt-like system of continuous tooth replacement, regulated by the maintenance of a stem cell population in the dental lamina, an essential structure for dental regeneration (Rasch et al., 2016, Fraser et al., 2020). This stem cell population is characterised by the expression of sex-determining region Y-related box 2 (*sox2*), an epithelial progenitor and stem cell marker (Martin et al., 2016, Juuri et al., 2013). Conversely, dermal denticles do not arise from a dental lamina and do not exhibit a continuous replacement mechanism. Instead, new denticles are thought to arise either as a result of growth of the body, denticle loss, or after wounding (Reif, 1978). Although a common odontode gene regulatory network (oGRN) (Fraser et al., 2010) appears to underpin the conserved development of both teeth and denticles, only teeth retain the ancestral gnathostome character of continuous successional tooth regeneration (Martin et al., 2016).

There are contrasting theories for the evolutionary origins of odontodes (Donoghue and Rucklin, 2016). One such theory suggests that external dermal odontodes arose first, before odontode-competent ectoderm subsequently migrated inside the oral cavity to form teeth (the ‘outside-in’ hypothesis). Conversely, it has been suggested that odontodes first arose inside the pharyngeal cavity, before migrating outwards to form dermal denticles (the ‘inside-out’ hypothesis) (Donoghue and Rucklin, 2016, Fraser et al., 2010) This uncertainty has arisen due to contrasting fossil evidence from early jawless vertebrates (Donoghue, 2002, Donoghue and Rücklin, 2016, Sire et al., 2009, Smith and Coates, 1998). Despite this ongoing debate, it is understood that odontodes can develop both inside and outside of the oral cavity (the ‘inside and out’ hypothesis), wherever conserved and co-expressed members of the underlying oGRN are present (Donoghue and Rücklin, 2016, Fraser et al., 2010). Importantly, studies examining the development of these units at the molecular and cellular levels have helped to resolve questions regarding the evolutionary origins of these skin appendages (Martin et al., 2016, Rasch et al., 2016).

Despite the structural similarities of elasmobranch dermal denticles and oral teeth, there is limited data directly comparing the embryonic development of these distinct structures. Here, we use immunohistochemistry (IHC) and in situ hybridisation (ISH) to characterise the cellular and molecular processes that underpin denticle development in the embryonic small-spotted catshark (*Scyliorhinus canicula*). The gene pathways examined here have been selected based on mammalian studies of dental development (Yu and Klein, 2020), although conservation of molecular signalling during elasmobranch tooth emergence has now been established (Rasch et al., 2016, Thiery et al., 2022, Rasch et al., 2020). Despite superficial differences in their form and function, we suggest that the early development and morphogenesis of shark odontodes is underpinned by the conserved expression patterns of a shared suite of developmental genes that comprise an oGRN, linking the molecular development of both teeth and scales.

## Methods

### Shark and chicken husbandry

The University of Sheffield is a licensed establishment under the Animals (Scientific Procedures) Act 1986. All animals were culled by approved methods cited under Schedule 1 to the Act. S. canicula embryos were purchased from North Wales Biologicals, UK, and raised in oxygenated artificial saltwater (Instant Ocean) at 16°C. Embryos were culled using MS-222 (Tricaine) at 300 mg/litre and fixed overnight in 4% paraformaldehyde in phosphate-buffered saline (PBS). Fertilized Bovan brown chicken eggs were purchased from Henry Stewart & Co., Norfolk, UK, incubated at 37.5°C, and fixed overnight in Carnoy’s solution. Following fixation, shark and chicken embryos were dehydrated through a graded series of PBS to ethanol (EtOH) and stored at -20°C.

### Scanning Electron Microscopy (SEM)

SEM was undertaken using a Hitachi TM3030Plus Benchtop SEM scanner at 15,000 V. Global brightness and contrast adjustments, and the addition of scalebars, was undertaken using Fiji (Schindelin et al., 2012).

### Alizarin red clear and staining

Fixed, dehydrated shark embryos were rehydrated into PBS and stained overnight in alizarin red in potassium hydroxide (KOH), as previously described (Cooper et al., 2017). Imaging was conducted using a Nikon SMZ15000 stereomicroscope, and scale bars were created using Fiji (Schindelin et al., 2012).

### Micro-Computed Tomography (Micro-CT)

Micro-CT scanning was undertaken using shark samples stained with 0.1% phosphotungstic acid (PTA) as previously described (Cooper et al., 2017), using an Xradia MicroXCT scanner at the Imaging and Analysis Centre of the Natural History Museum (London, UK). Rendering was undertaken using the 3D volume exploration tool Drishti (www.github.com/nci/drishti).

### Immunohistochemistry (IHC)

IHC of paraffin sections was undertaken as previously described (Rasch et al., 2016). Sections were imaged with an Olympus BX51 Upright Compound Microscope and Olympus DP71 Universal digital camera attachment. Fiji was used to globally adjust brightness and contrast and to add scale bars (Schindelin et al., 2012).

### In situ hybridisation

The design of digoxigenin-labelled antisense riboprobes and subsequent whole mount in situ hybridization was undertaken as previously described (Cooper et al., 2017, Cooper et al., 2018). Riboprobes were designed using partial skate (*Leucoraja erinacea*) and catshark (*S. canicula* or *Scyliorhinus torazame*) EST assemblies (Wyffels et al., 2014) (SkateBase, skatebase.org), and the Vertebrate TimeCapsule (VTcap, transcriptome.cdb.riken.go.jp/vtcap). The riboprobes were cloned from *S. canicula* cDNA using the primer sequences shown in Table 1. Sections were imaged with an Olympus BX51 Upright Compound Microscope and Olympus DP71 Universal digital camera attachment. Whole mount samples were imaged using a Nikon SMZ15000 stereomicroscope. Fiji was used to globally adjust brightness and contrast and to add scale bars (Schindelin et al., 2012). The riboprobes were cloned from S. canicula cDNA using the following primer sequences, sequence databases and published GenBank accession numbers:

**Table 1:**
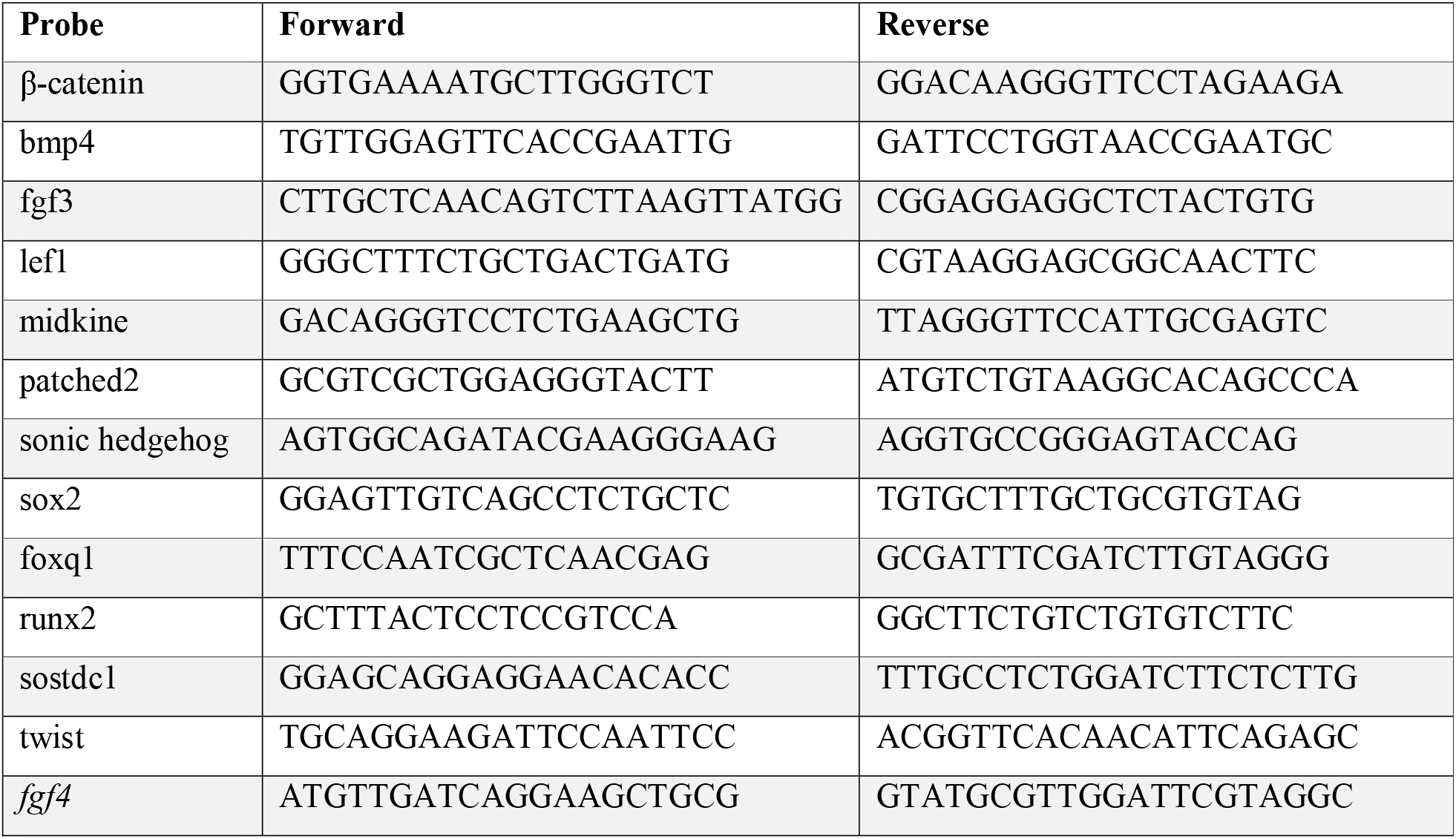
Primer sequences for *S. canicula* riboprobes

### Whole mount in situ hybridisation

The design of digoxigenin-labelled antisense riboprobes and subsequent whole mount in situ hybridization was undertaken as previously described (Cooper et al., 2017, Cooper et al., 2018). Samples were imaged using a Nikon SMZ15000 stereomicroscope. Fiji was used to globally adjust brightness and contrast and to add scale bars (Schindelin et al., 2012).

### Whole mount immunofluorescence

Samples were rehydrated from EtOH through a graded series of PBS with 0.5% Triton (PBST) and treated with 10μg/ml proteinase k for 20 minutes. Samples were then incubated in 5% goat serum with 1% bovine serum albumin in PBS for the blocking stage. Primary antibody staining took place for 2 days at 4°C, using both Anti-SHH (AV44235, Sigma-Aldrich) and Anti-PCNA (ab29, Abcam) at a concentration of 1:500. Incubation in the secondary antibody was performed under the same conditions, using goat anti-mouse Alexa Fluor 488 and goat anti-rabbit Alexa Fluor 647 (Thermo Fisher) respectively. Samples were counterstained with DAPI before imaging with a Zeiss LMS 880 with Airyscan. Images shown in Figure 7 were composed using the standard deviation projection of a Z-series in Fiji (Schindelin et al., 2012).

## Results

### Odontode diversity in the small-spotted catshark

First, we use a combination of scanning electron microscopy (SEM) and micro-computed tomography (micro-CT), to explore odontode diversity in the small-spotted catshark (*S. canicula*) (Fig. 1). *S. canicula* exhibits various different odontode types that can be broadly organised into four distinct categories: (a) transient denticles of the caudal tail, (b) enlarged denticles of the dorsal trunk, (c) adult-type body denticles, which exhibit multiple forms, and (d) multicuspid oral teeth (Cooper et al., 2017, Ballard et al., 1993).

**Figure 1:**
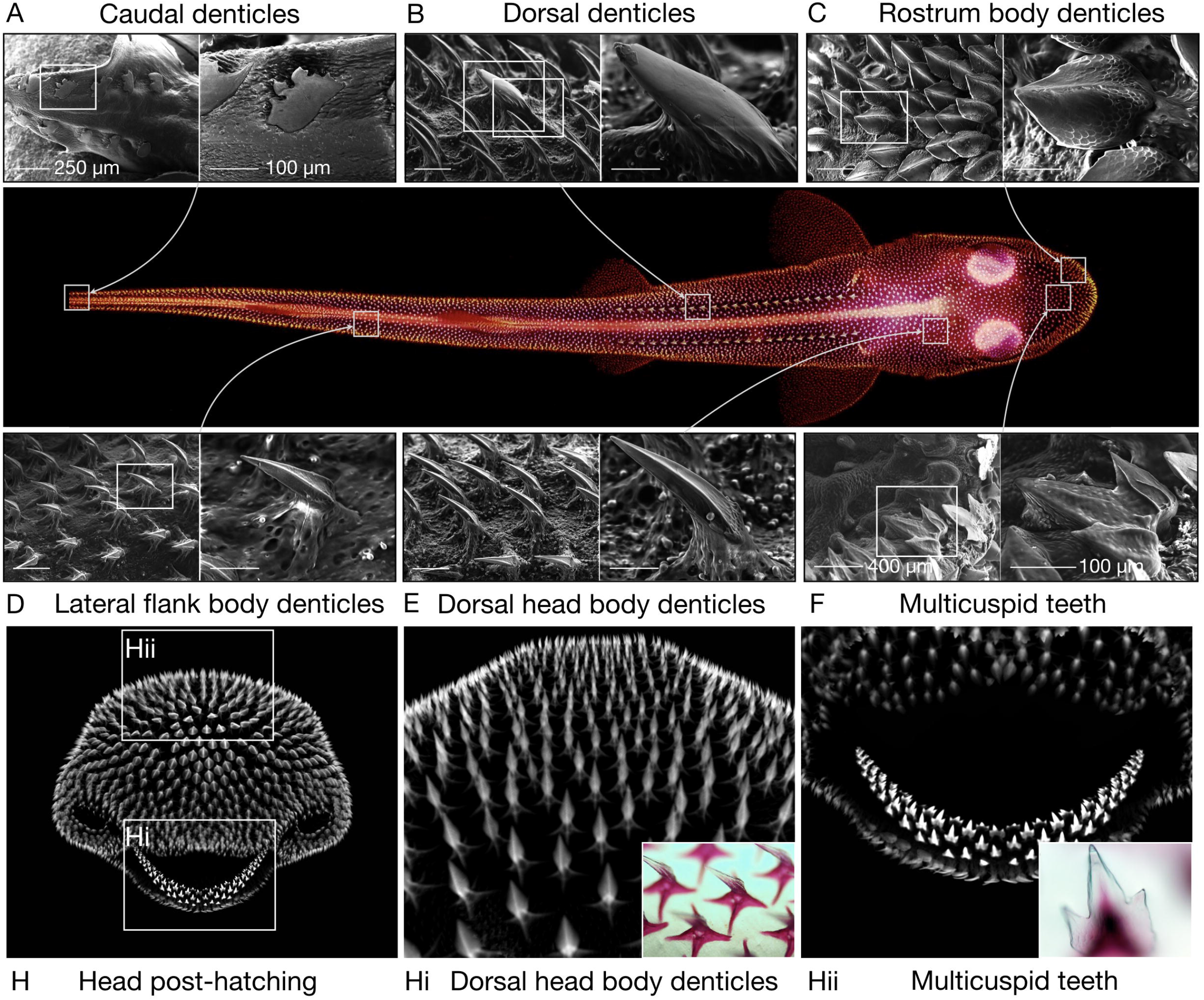
Odontode diversity of the small-spotted catshark. Scanning Electron Microscopy (SEM) is used to reveal odontode diversity in the shark. Caudal denticles are the first odontode type to emerge, in two lateral and dorsal rows, either side of the tip of the tail (A). These flattened units possess irregular posterior facing cusps, and an ancient dentine type associated with sharks from the Silurian and Ordovician (Johanson et al., 2008). Two dorsolateral rows of enlarged denticles with rounded cusps next emerge on the dorsal trunk (B). These dorsal rows initiate the wider propagation of body denticles (C-E) (Cooper et al., 2018), which exhibit diverse forms whilst consistently displaying a single cusp. Multicuspid teeth emerge close to 110 dpf, initially with a tricuspid morphology (Fig. F), although cusp number increases with tooth replacement (Thiery et al., 2022). This odontode diversity is also shown with micro-CT of the shark head (H), in which diverse body denticle types (Hi) and multiple generations of multi-cuspid teeth (Hii) are visible. Insets in panels Hi and Hii show alizarin red stained denticles and teeth, respectively. The whole hatchling shark is an alizarin red preparation.

Caudal denticles of the posterior tail are the first odontode type to emerge, arising between 52-60 days post fertilisation (dpf), in two dorsal and ventral rows positioned laterally on either side of the tail fin tip (Fig. 1A) (Johanson et al., 2007). Caudal denticles are placode-derived skin appendages (Cooper et al., 2017) with, typically, between 9 and 13 units forming on either dorsal row, and between 5 and 10 units forming on either lateral row (Ballard et al., 1993). These flattened units develop sequentially from posterior to anterior, approximately equidistant from one another, and exhibit highly irregular, posterior facing cusps. Caudal denticles contain an ancient dentine type constructed from tubules that exhibit a distinct branching pattern associated with the earliest known sharks from the Ordovician and Silurian. Therefore, these denticles are considered an ancestral odontode type (Johanson et al., 2008). Interestingly, these are transient units that are lost close to the time of hatching, when adult type body denticles arise to occupy their positions.

Two dorsolateral rows of enlarged denticles on the dorsal trunk are the second odontode type to emerge, between 60-80 dpf (Fig. 1B) (Ballard et al., 1993, Enault et al., 2016). These units lack distinct ridges and have a rounded posterior-facing cusp. They are subsumed into general scalation shortly after hatching (Martin et al., 2016). Dorsal denticles act as initiator rows, triggering the subsequent emergence of body denticles via a Turing-like reaction-diffusion system (Cooper et al., 2018), comparable to the patterning of avian feathers and scales (Jung et al., 1998, Cooper et al., 2019).

Body denticles are the last denticle type to emerge, propagating across the entire body surface at approximately 100 dpf (Fig. 1C-E, H) (Cooper et al., 2018). They exhibit dramatic variation in their morphology (Gabler-Smith et al., 2021), ranging from the petal-like denticles of the rostrum (Fig. 1C) to the sharp, protruding denticles of the dorsal head surface (Fig. 1D, H) and lateral flank (Fig. 1E) which exhibit distinct ridges, likely associated with hydrodynamic drag reduction (Oeffner and Lauder, 2012, Wen et al., 2015). Although body denticles exhibit substantial morphological variation, they typically display a single posterior facing cusp (Fig. 1C-E, H). Conversely, oral teeth, which begin to emerge at approximately 110 dpf, are initially tricuspid (Fig. 1F, Hii). However, after multiple rounds of tooth replacement, five or more cusps can be observed (Thiery et al., 2022). Overall, we report a vast diversity of odontode morphologies within a single shark species.

### Cell proliferation dynamics through dorsal denticle emergence

Next, we use IHC for proliferating cell nuclear antigen (PCNA) to understand the cellular and tissue layer proliferative processes involved in dorsal denticle emergence and morphogenesis in the embryonic shark (Fig. 2).

**Figure 2:**
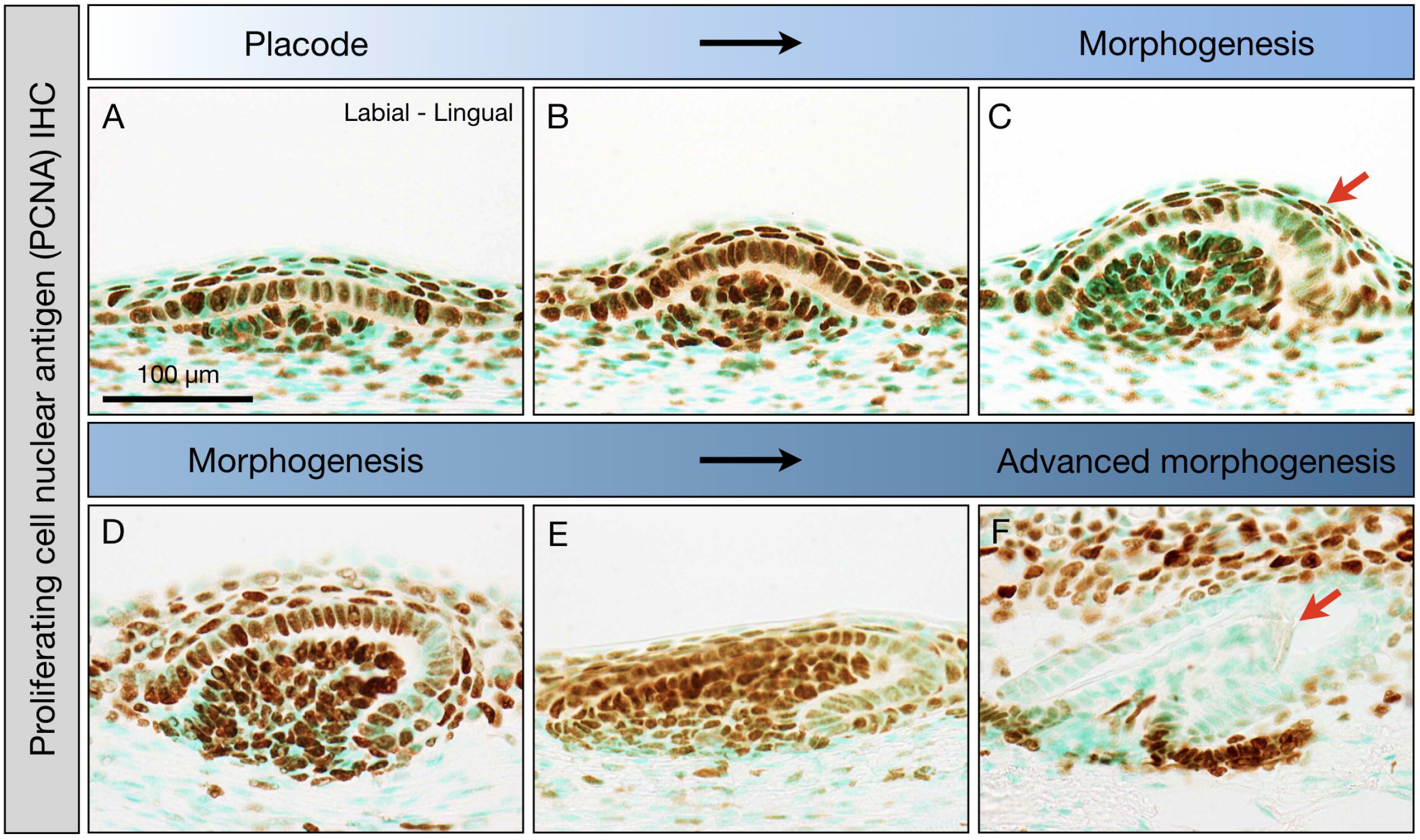
Cell proliferation during dorsal denticle emergence. IHC for PCNA is used to examine proliferative processes involved in dorsal denticle development. Denticle placode initiation is marked by a local epithelial thickening, associated with an underlying dermal condensation (A). Continued cell proliferation underpins the evagination of the epidermis and condensing dermis (B). Onset of morphogenesis is marked by asymmetric outgrowth, accompanied by reduced cell proliferation in the distal epithelial tip, in contrast to the adjoining epithelium and underlying mesenchyme (C, red arrow). This reduced proliferation continues, focal to the epithelial tip (D-E). Following advanced morphogenesis (E), cell proliferation in both the epithelium and mesenchyme is negligible, marked by a total lack of PCNA immunoreactivity of the denticle unit (F). Secreted mineralised tissue can be seen in the papilla (F, red arrow) suggesting terminal differentiation of cells to both ameloblasts and odontoblasts.

Denticle placode initiation is marked by localised epithelial and mesenchymal cell proliferation. Basal epithelial cells become columnar, producing a localised thickening of the epidermis, associated with an underlying dermal cellular condensation (Fig. 2A). Early morphogenesis involves the evagination of the placode (Fig. 2B), followed by onset of growth polarity and a reduction in cell proliferation in the distal epithelial tip, shown by reduced PCNA immunoreactivity (Fig 2C, red arrow). Subsequent morphogenesis is further accompanied by progressive enclosure of the proliferating mesenchymal compartment (Fig. 2D). Interestingly, this non-proliferative region of the denticle cusp is indicative of a signalling centre comparable to the enamel knot, a conserved mediator of tooth cusp formation (Thiery et al., 2022, Jernvall and Thesleff, 2012, Vaahtokari et al., 1996). As polarised growth continues, a reduction of PCNA immunoreactivity in the epithelial tip is maintained (Fig. 2E). Throughout advancing morphogenesis, a total reduction in PCNA immunoreactivity in both the epithelium and mesenchyme implies terminal differentiation of cells to ameloblasts and odontoblasts, respectively (Fig. 2F). Corresponding matrix deposition in the papilla is also apparent (Fig. 2F, red arrow). Overall, these results suggest comparable proliferative developmental processes between both dermal denticle and oral tooth cusp formation (Thiery et al., 2022).

### Conserved patterns of tooth-associated gene expression are deployed throughout denticle development

To explore the potential deployment of a shared genetic toolkit common to all odontodes, the expression patterns of genes representing several signalling pathways were investigated during body denticle development using *IS*H (Fig. 3). We examined the expression of genes known to be involved in both epithelial and mesenchymal contributions to mammalian and non-mammalian tooth development (Jernvall and Thesleff, 2000).

**Figure 3:**
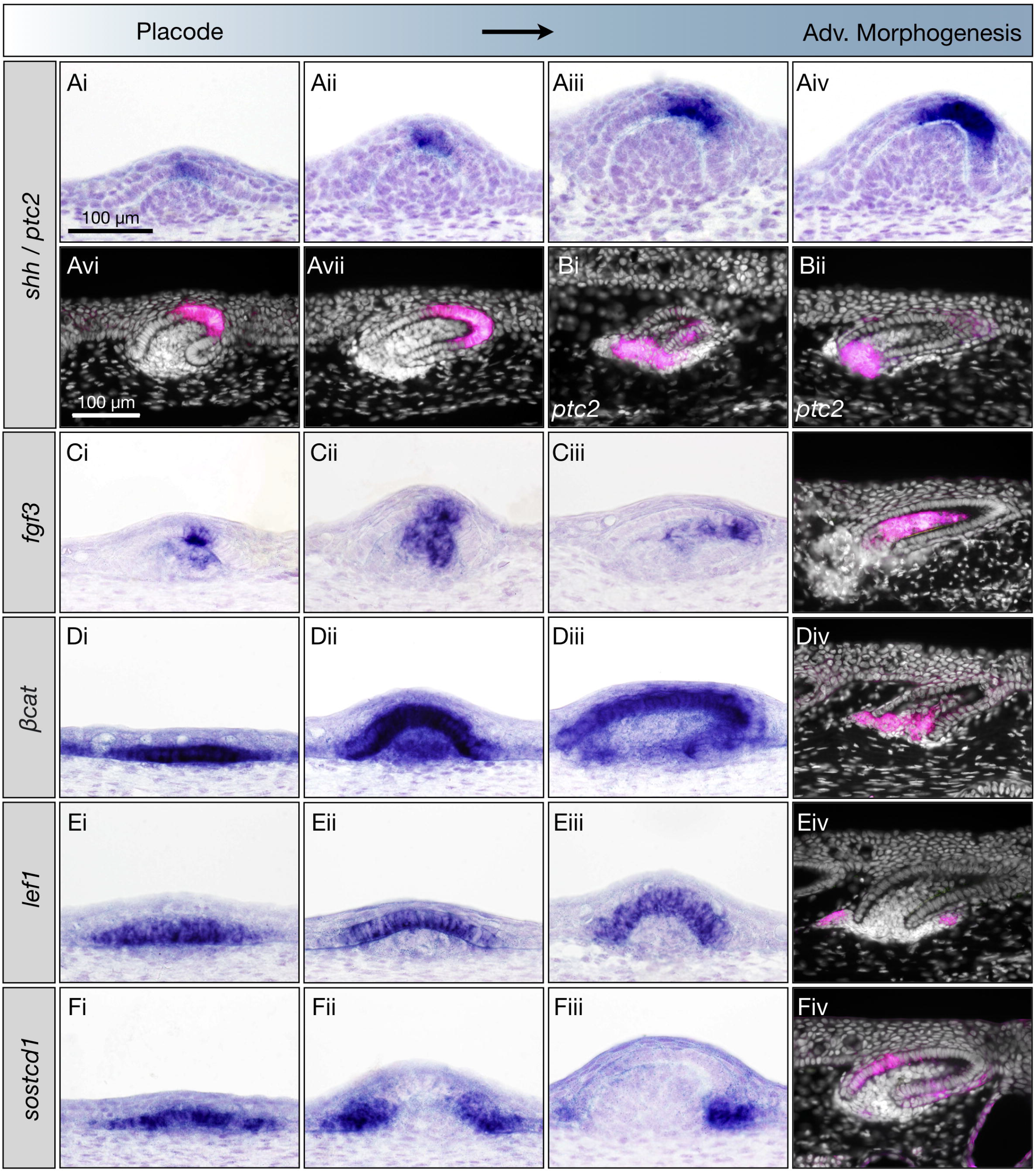
Gene expression patterns associated with shark denticle development. *shh* expression is observed in early developing denticles (A), increasing in intensity in the outgrowing cusp during morphogenesis and advanced morphogenesis (Ai-AVii). Expression of the shh receptor, *ptch2*, is also noted in the basal mesenchyme directly underlying the papilla during morphogenesis (Bi – Bii). *fgf3* is first weakly expressed in the medial mesenchyme directly underlying the epithelial basal membrane (Ci), before becoming restricted to an asymmetric region of the epithelium (Cii-Ciii). Diffuse staining of β*cat* is observed in regions of the basal epithelium, marking placode initiation (Di), and remains present during subsequent morphogenesis (Dii – Diii). By advanced morphogenesis, β*cat* is restricted to the basal mesenchyme of the papilla (Div). During placode initiation, *lef1* is expressed in the basal epithelium (Ei), and its expression is maintained throughout subsequent morphogenesis, before becoming restricted primarily to two bilateral regions of the basal mesenchyme adjacent to the papilla (Eii - Eiv). We also observed expression of *sostdc1* in the basal epithelium of each placode forming unit (Fi), with expression becoming progressively restricted to the peripheral epithelium during subsequent morphogenesis, leaving a *sostcd1*-ve medial region (Fii-Fiv). False coloured (magenta) images are counterstained with DAPI.

Sonic hedgehog (*shh*) is a well-known facilitator of epithelial appendage patterning and development (Busby et al., 2020, Chiang et al., 1999, Chuong et al., 2000), and is specifically expressed at several key stages of tooth development (Cho et al., 2011, Smith et al., 2009, St-Jacques et al., 1998). It has also been observed in the developing caudal denticles of the catshark (Johanson et al., 2008, Cooper et al., 2017). Here, we observe *shh* in early developing body denticle placodes, and subsequently in the apical cells (central inner dental epithelial cells; equivalent to the Apical Epithelial Knot (AEK) described in the shark tooth (Thiery et al., 2022) of the early denticle bud (Fig. 3Ai-Aii). As the bud advances through morphogenesis, *shh* expression becomes restricted to a distinct region of the epithelium, showing an initial polarity bias towards the posterior aspect of the apex (Fig. 3Aiii-Aiv). Throughout advanced morphogenesis, *shh* expression persists in the AEK, localising to the polarised distal epithelial tip and spreading to neighbouring cells at the cusp apex (Fig. 3Aiv-vii). *shh* signals to target cells via its receptor, patched 2 (*ptch2*) (Ingham and McMahon, 2001). Here, *ptch2* expression is present within the basal mesenchyme of the denticle papilla (Fig. 3Bi-Bii), and is weakly expressed within cells surrounding the *shh*+ apex epithelium.

Fibroblast growth factor 3 (*Fgf3*) is a highly conserved member of the fibroblast growth factor family of signalling molecules, expressed during tooth cusp, hair follicle, feather bud and caudal denticle development (Bei and Maas, 1998, Cooper et al., 2017, Fraser et al., 2013, Jackman et al., 2004, Kettunen et al., 2000, Mandler and Neubuser, 2004, Rosenquist and Martin, 1996). During denticle development, *fgf3* is first detected at the placode stage during denticle development, localised in both the epithelium and underlying medial mesenchyme (Fig. 3Ci). As denticle morphogenesis progresses, *fgf3* expression spreads to encompass more of the papillary mesenchyme, accompanied by a marked increase in expression concentrated in the apex of the denticle epithelium (Fig. 3Cii). Throughout subsequent morphogenesis, the polarised epithelial-mesenchymal expression pattern of *fgf3* progressively increases (Fig. 3Ciii), before finally becoming restricted entirely to the mesenchymal papilla (Fig. 3Civ).

The intracellular signal transducer of the Wnt signalling pathway, β-catenin (β*cat*), is required for the initiation and morphogenesis of hair follicles, feather buds and teeth (Noramly et al., 1999, Chen et al., 2012, Jarvinen et al., 2006, Millar et al., 1999). Developing denticles show intense β*cat* expression associated with all developmental stages, from the early placode to later stages of morphogenesis (Fig. 3D). Expression is first restricted to the basal epithelium of each placode-forming unit (Fig. 3Di). Throughout subsequent stages of bud formation, this epithelial expression is sustained, further spreading to the underlying condensed mesenchyme and the developing papilla (Fig. 3Dii-iii). By advanced morphogenesis, β*cat* expression is completely absent from the epithelium, restricted instead to the basal mesenchyme of the denticle papilla (Fig. 3Div).

Throughout activation of Wnt signalling, nuclear β*cat* activates target genes by binding with lymphoid enhancing factor 1 (*lef1*), which is also prominently expressed during both tooth and feather development (Seidensticker and Behrens, 2000, Chen et al., 2009, Handrigan and Richman, 2010). During development of denticle primordia, *lef1* is initially expressed in a similar pattern to β*cat*, marking individual placodes via expression in the basal epithelium (Fig. 3Ei-ii). However, throughout subsequent outgrowth *lef1* becomes restricted to the epithelial cells of the denticle placode (Fig. 3Eii-Eiii). By advanced morphogenesis, *lef1* expression is primarily restricted to two bilateral regions of the basal mesenchyme adjacent to the papilla (Fig. 3Eiv).

The secreted sclerostin domain-containing protein 1 (Sostcd1/Ectodin/Wise) interacts with bone morphogenic protein (BMP), Wnt, FGF and Shh signalling to regulate the spatial patterning and morphogenesis of teeth, and the development of other epithelial appendages (Ahn et al., 2010, Cho et al., 2011, Munne et al., 2009, Mou et al., 2011). In shark denticle development, *sostdc1* is expressed in the basal epithelium of the placode (Fig. 3Fi). During early growth, expression shifts bilaterally to the peripheral epithelium (equivalent to the outer dental epithelium (ODE) in teeth), leaving a central apical region devoid of expression (Fig. 3Fii-Fiii). By advanced morphogenesis, *sostdc1* becomes predominantly restricted to the posterior in-fold of the epithelium towards the base of the denticle cusp (Fig. 3Fiv).

The heparin-binding growth factor Midkine (*mk*) regulates various aspects of cell growth and differentiation and is also expressed throughout different stages of tooth development (Mitsiadis et al., 2008, Mitsiadis et al., 1995, Park et al., 2020). Throughout denticle emergence, *mk* is first observed in the thickened epithelium of denticle placodes, with expression also noted in the underlying mesenchyme (Fig. 4Ai). Mesenchymal expression of *mk* subsequently expands to encompass the entire dental papilla (Fig. 4Aii). During later morphogenesis, expression in the distal epithelial tip increases (Fig. 4Aiii). Subsequent advanced morphogenesis is marked by maintenance of papillary expression, and a reduction in expression at the polarised epithelial tip (Fig. 4Aiv).

**Figure 4:**
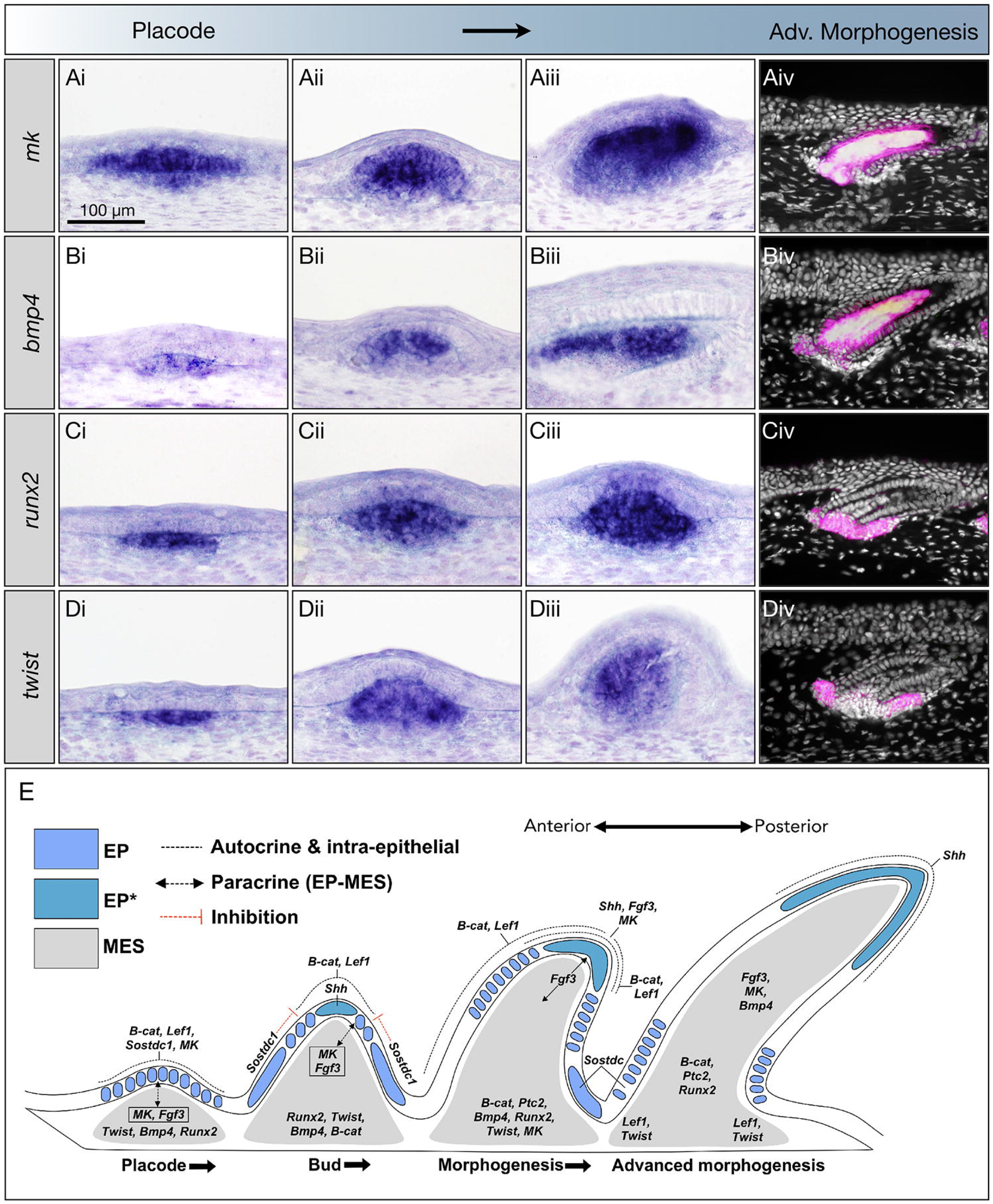
Dermal gene expression patterns associated with shark denticle development. *mk* is strongly expressed in both the epithelium and underlying mesenchyme, from placode initiation through to advanced morphogenesis (Ai-Aiv). We also observe mesenchymal expression of *bmp4* (Bi-Biv), *runx2* (Ci-Civ) and *twist* (Di-Div) throughout dermal denticle development, from placode initiation to advanced morphogenesis. During advanced morphogenesis, *bmp4* expression highlights the entire dermal papilla (Biv), whereas *runx2* and *twist* become restricted to the base of the outgrowing unit (Civ, Div). False coloured images are counterstained with DAPI. We also present a putative denticle GRN (E). We propose that molecular signalling cascades regulate denticle development from placode stage to advanced morphogenesis (from left to right; Anterior to Posterior), through expression of activators, inhibitors, polarising growth signals and differentiation factors. β*cat* and *lef1* are proposed to positively regulate cell proliferation during development (Jarvinen et al., 2006, Noramly et al., 1999), while inhibitors such as *sostdc1* and *bmp4* can precisely delineate these expression domains (Cho et al., 2011, Noramly and Morgan, 1998, Mou et al., 2011). Dynamic expression of *fgf3* and *mk* between the epithelium and mesenchyme suggests roles in mediating inductive tissue interactions (Mitsiadis et al., 2008, Kettunen et al., 2000). In association with β*cat*, progressive localisation of *shh, fgf3* and *mk* to the non-proliferative epithelial tip infers roles as polarizing factors, guiding unidirectional growth, and regulating subsequent cusp formation. This signaling centre is proliferatively (Fig. 2) and molecularly comparable to the mammalian enamel knot (Vaahtokari et al., 1996), which is also implicated in the development of dental cusps in elasmobranch teeth (Thiery et al., 2022). *Twist* is also proposed to act in accordance with its conserved role as a negative regulator of *runx2*, in advance of its function inducing cell differentiation for matrix deposition (Bialek et al., 2004). EP, epithelium; EP*, epithelial tip; MES, mesenchyme; EP-MES, epithelial-mesenchymal.

BMPs also regulate various aspects of tooth, feather, and hair follicle development by mediating epithelial-mesenchymal interactions (Åberg et al., 1997, Mou et al., 2011, Noramly and Morgan, 1998, Vainio et al., 1993). In particular, mesenchymal expression of *Bmp4* is highly conserved during skin appendage placode formation (Cooper et al., 2017, Di-Poï and Milinkovitch, 2016). In denticle development, *bmp4* is initially weakly expressed in the placode mesenchyme (Fig. 4Bi), with expression subsequently increasing in intensity as denticle outgrowth continues (Fig. 4Bii). Expression remains restricted to the epithelium as morphogenesis progresses, marking the dermal papilla of the developing denticle (Fig. 4Biii). Later, this pattern is accompanied by epithelial expression at the cervical-loop-like basal regions of the denticle, equivalent to the cells at the junction between the inner and outer dental epithelium of developing teeth (IDE/ODE; Fig. 4Biv).

Of the Runt-related (Runx) family of transcription factors, *Runx2* regulates bone calcification by controlling the proliferation and differentiation of cells committed to osteoblastic lineages (Camilleri and McDonald, 2006). The deeply conserved odontogenic role of *runx2* has previously been shown by expression in both teeth and denticles of the catshark, implying its co-option from a common developmental module to allow the evolution of odontodes (Hecht et al., 2008). Here we further investigate the expression patterns of *runx2* at various stages of denticle development. Expression is first detected in the mesenchyme underlying the early denticle placode (Fig. 4Ci). During subsequent bud outgrowth, this mesenchymal expression is maintained (Fig. 4Cii), however, expression subsequently spreads from the mesenchymal compartment into the directly overlying medial/apical epithelium (Fig. 4Ciii). In later stages of morphogenesis, expression becomes restricted to the basal mesenchyme of the advancing papilla and the surrounding deeply invaginated epithelial loops (equivalent to the dental cervical-loop-like regions; Fig. 4Civ).

The Twist transcription factor is also associated with bone development, via regulation of osteoblastic cell activity (Murray et al., 1992, Rice et al., 2000). Here, *twist* is first detected in the mesenchyme (Fig. 4Di), in a comparable pattern to *runx2* (Fig. 4Ci). Further similarities with *runx2* are observed throughout placode outgrowth (Fig. 4Dii). During morphogenesis, however, *twist* shows restriction to the anterior aspect of the mesenchymal papilla, leaving an apparent negative region of posterior mesenchyme (Fig. 4Diii). By advanced stages of morphogenesis, *twist* becomes progressively restricted to the bilateral periphery of the basal mesenchyme of the papilla (Fig. 4Div) surrounding the epithelial cervical loop-like regions of the denticle (ODE/IDEO equivalent cells).

The expression data presented here represents an initial framework for a hypothetical denticle gene regulatory network (dGRN) model, which shows broad conservation with the oral dentition, therefore expanding our knowledge of the conserved and ancient oGRN (Fig. 4E) (Martin et al., 2016, Rasch et al., 2016, Rasch et al., 2020). We observe the expression of several well-known tooth-related genes in an almost equivalent manner to early dental morphogenesis (Fig. 3, Fig. 4). We suggest that, whether odontodes first appeared in the oral/pharyngeal cavity or within the ectodermal epidermis, the oGRN likely emerged to form tooth-like structures prior to the evolution of true vertebrate teeth, and the mechanisms of tooth regeneration from within an active dental lamina likely evolved later.

### Shared gene expression patterns throughout oral tooth and dermal denticle development

Next, we undertook section *IS*H of the shark lower jaw, which contains both denticles and teeth. This enabled us to examine patterns of simultaneous gene expression in these distinct odontode types (Fig. 5).

**Figure 5:**
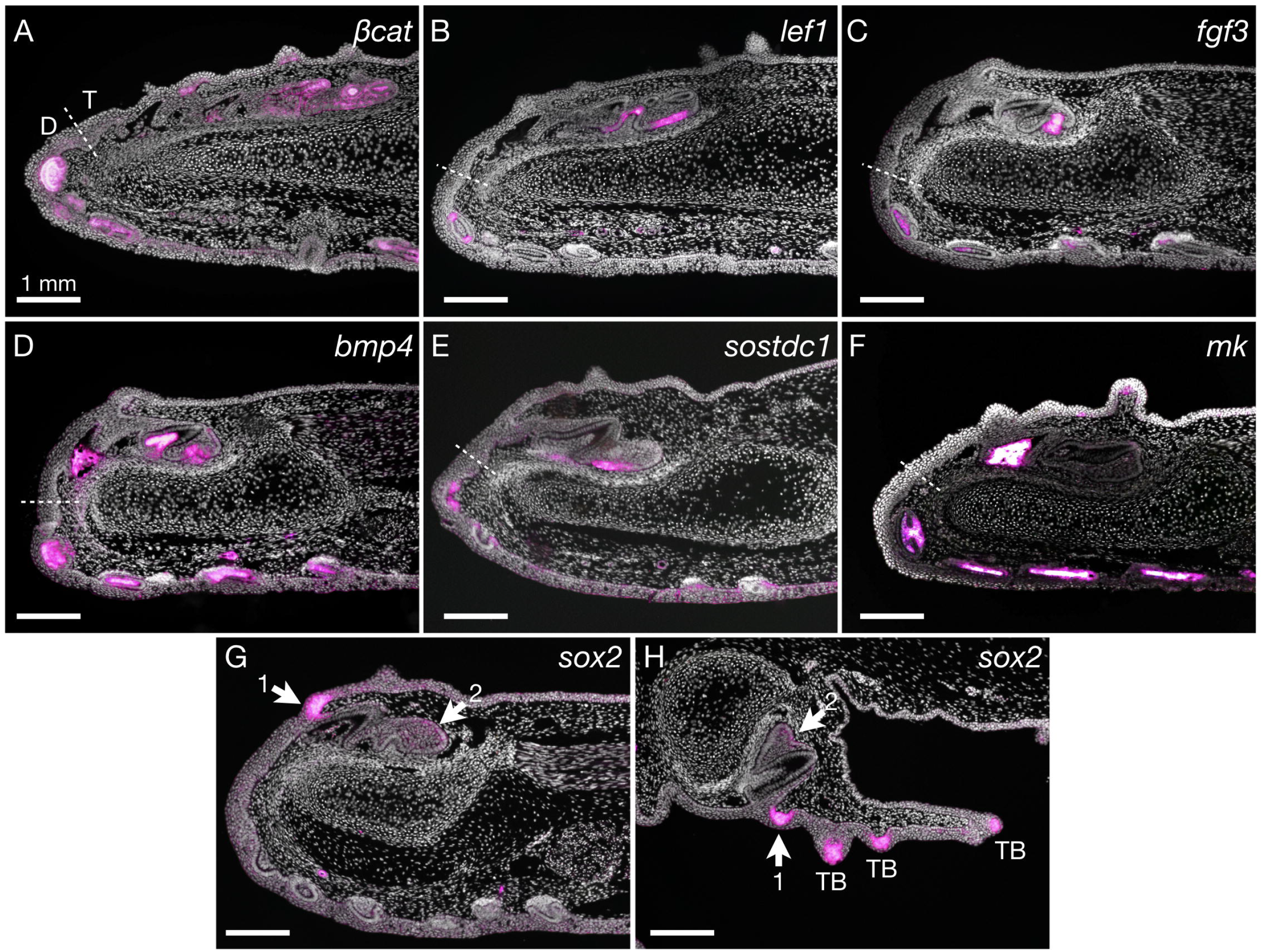
Gene expression patterns during development of both the regenerative shark dentition and the skin denticles. In both dermal denticles and oral teeth of the lower jaw at ∼110 dpf (sagittal plane of section), simultaneously expressed genes imply the deployment of a common oGRN. In denticle and tooth development, commonly expressed β*cat, lef1, fgf3, bmp4, sostdc1* and *mk* (A-F, respectively) define these key similarities, shown by their expression in both oral teeth and dermal denticles. Teeth (T) and denticles (D) are further demarcated by a putative boundary zone (dotted line) between the oral and dermal epithelia. Differential expression of *sox2*, confined to the dental lamina (G-H), provides a key difference between denticles and teeth. In the oral epithelium, *sox2* shows clear associations with developing teeth, where its expression marks a putative dental stem cell niche (G, arrow 1) linked with the successional lamina via a continuous epithelial stripe (H, arrow 2). Additionally, *sox2* marks regenerative taste buds (H, TB). This indicates a difference between inner (oral) and outer (dermal) epithelia, as defined by a *sox2+* dental lamina that facilitates continuous tooth replacement, in contrast to *sox2-* dermal denticles, which do not exhibit a system of continuous replacement. Orientation: Anterior to the left; sagittal plane of section.

In concurrence with our previous results (Fig. 3, Fig. 4), we observe conservation of gene expression patterns between both oral teeth and dermal denticles, including those of β*cat, lef1, fgf3, bmp4, sostdc1* and *mk* (Fig. 5A-F). However, although the transcription factor *sox2* is expressed in the developing dentition, it remains absent from the emerging denticles (Fig. 5G) (Martin et al., 2016). We also note expression of *sox2* in the developing regenerative taste buds (Fig. 5H). This transcription factor has a conserved role related to the maintenance of the stem cell niche, essential for vertebrate tooth initiation and regeneration (Juuri et al., 2013, Martin et al., 2016, Fraser et al., 2020). This lack of expression in denticles may explain the difference in regenerative potential between these distinct odontode types.

### Tooth-associated gene expression patterns are deployed throughout development of the shark sensory system

Various epithelial sensory structures are located within close proximity of developing body denticles. Therefore, we next sought to compare the expression of several genes between emerging body denticles and regenerative taste buds, pit organs and functional Ampullae of Lorenzini in the external skin of the epithelium (Fig. 6).

**Figure 6:**
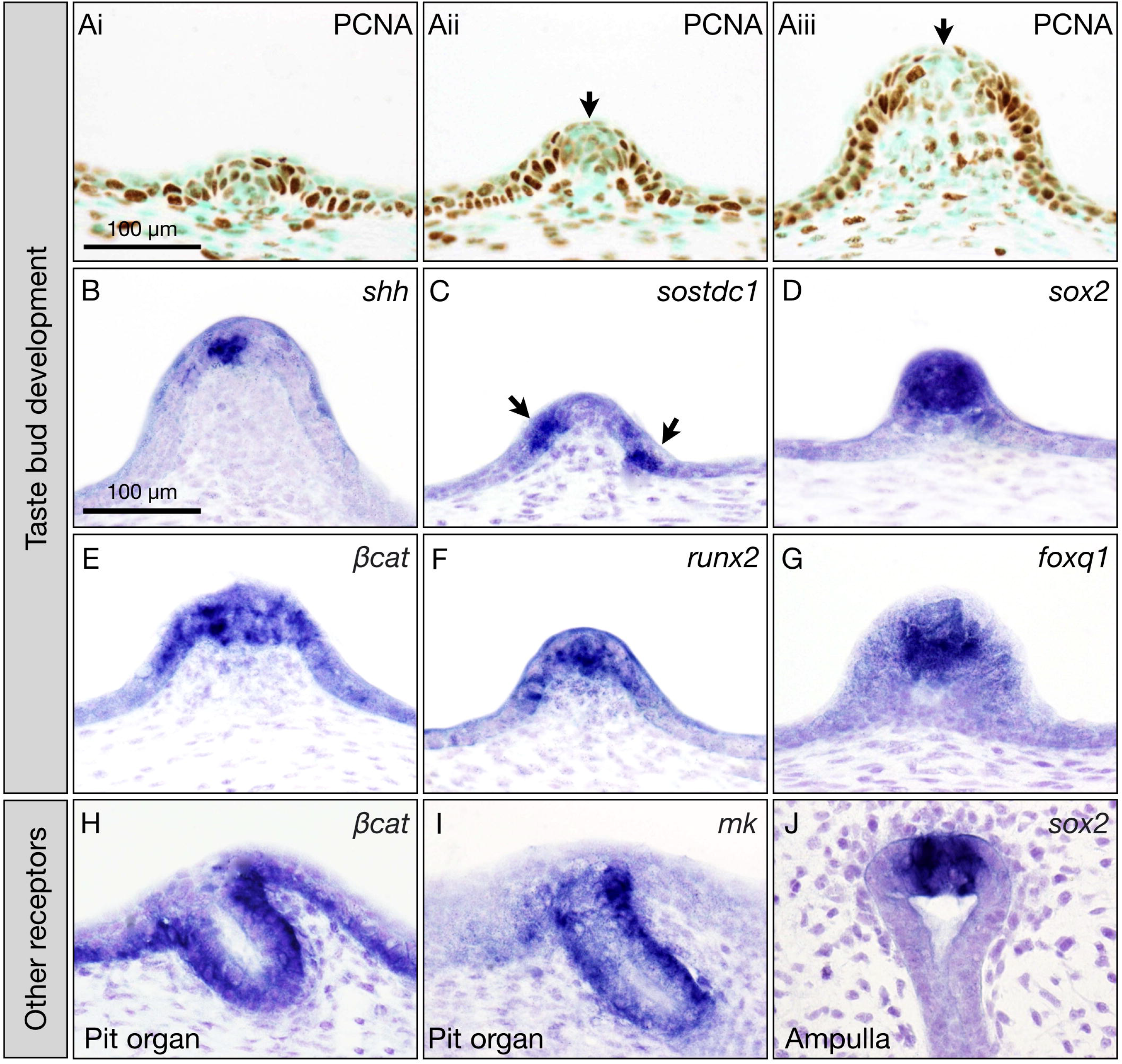
Cell proliferation and conserved gene expression patterns associated with the development of sensory receptors in the shark. PCNA immunoreactivity reveals that taste bud papillae arise through controlled proliferation of the epithelium from the placode stage (Ai) to later morphogenesis (Aii-Aiii). Reduced proliferation in the papillae tip is also noted here (Aii-Aiii, black arrow). At similar stages, *shh* is expressed in the evaginating epidermis (B). At preceding stages, *sostdc1* is expressed bilaterally in the periphery of early taste bud papillae (C, arrows). Additional markers, including *sox2*, β*cat, runx2*, and *foxq1* (D-G) are expressed in the epithelium at various developmental stages. Additionally, β*cat* and *mk* are also expressed in developing pit organs, and *sox2* can be observed in the sensory cells of the Ampullae of Lorenzini.

PCNA immunoreactivity of taste bud paraffin sections reveals outgrowth of a highly proliferative epithelial thickening (Fig. 6Ai). Interestingly, we also observe a medial zone of reduced proliferation throughout taste bud morphogenesis (Fig. 6Aii-Aiii), indicative of comparable proliferative growth to dorsal denticles (Fig. 2). We also note similarities in specific gene expression patterns during taste bud morphogenesis. For example, *shh* expression is concentrated in the apical tip of the taste epithelium (Fig. 6B), and *sostdc1* is restricted to lateral epithelial cells surrounding the apex of each taste bud (Fig. 6C). *sox2* contributes to the development of various sensory structures (Castillo-Azofeifa et al., 2018, Martin et al., 2016, Okubo et al., 2006), and here we observe its expression in developing taste buds (Fig. 6C) and the Ampullae of Lorenzini sensory neuromast (Fig. 6J). Conserved expression of β*cat* and *runx2* are also observed (Fig. 6E-F). Additionally, we observe expression of Fox gene family member, *foxq1*, a known inducer of epithelial differentiation, during shark taste bud development (Fig. 6G) (Feuerborn et al., 2011). β*cat* and *mk* are also present during pit organ development (Fig. 6H-I). Overall, we observe broad conservation of tooth-related gene expression patterns during the early development of the shark sensory system.

### Conserved gene expression patterns are deployed during the early development of both shark denticles and avian feathers

Having observed similarities between expression patterns of tooth-related genes throughout development of both denticles and the shark sensory system, we next sought to examine whether this conservation extends beyond sharks, to encompass the skin appendages of other species. Previous research has demonstrated that the anatomical placode constitutes a common foundation from which diverse skin appendages are derived across phylogenetically distinct vertebrate groups (Di-Poï and Milinkovitch, 2016, Cooper et al., 2017, Harris et al., 2008). Using whole mount *IS*H, we compared expression patterns of conserved developmental genes during skin appendage development of both the shark and the chicken embryo (Fig 7).

**Figure 7:**
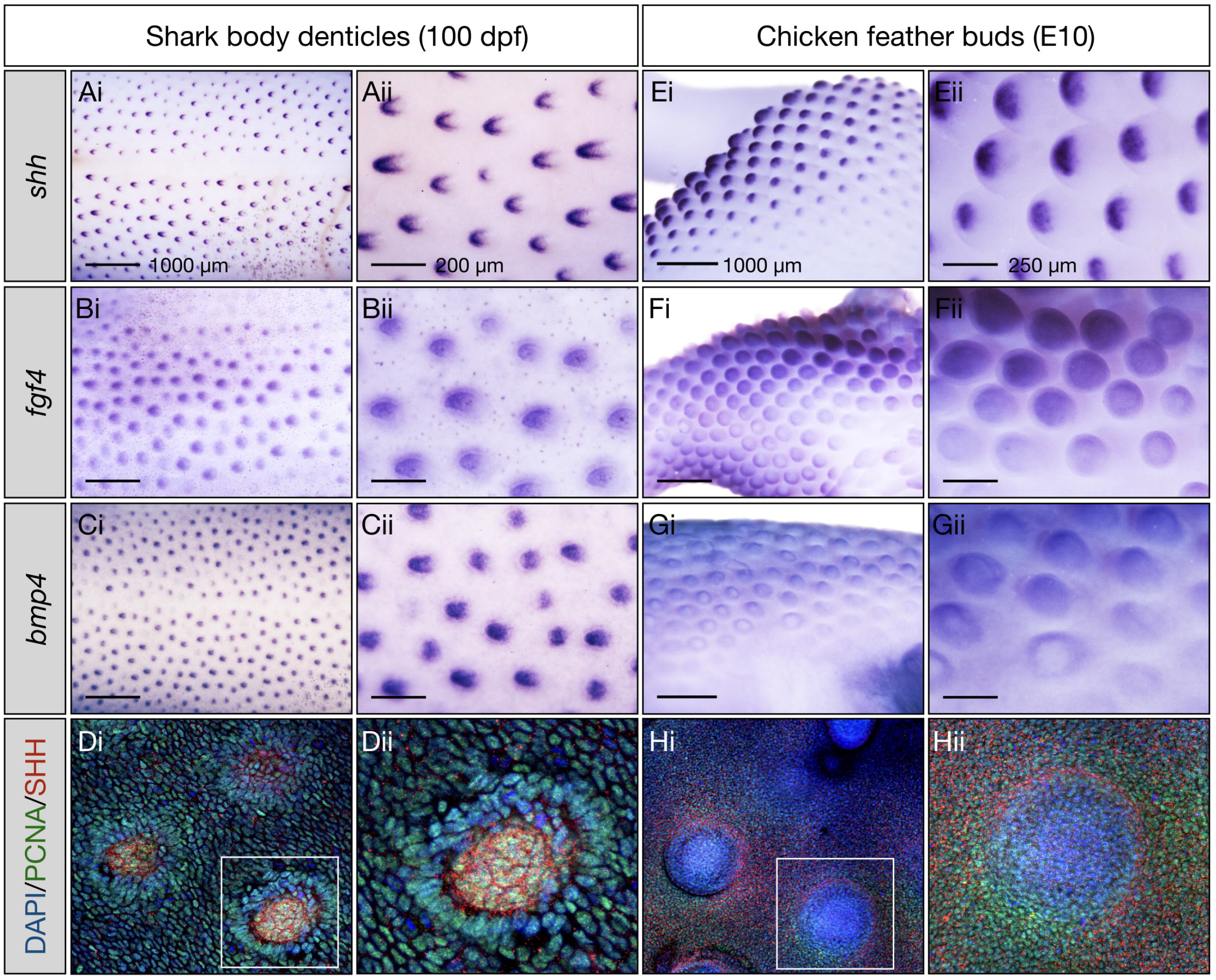
Conserved gene expression patterns during early shark denticle and avian feather development. The development of both shark denticles and chicken feathers is compared, using whole mount ISH. The early development of shark body denticles is observed at 100 dpf (A-D). Epithelial expression of *shh* is observed in the posterior cusp of developing denticles (A), whereas *fgf4* and *bmp4* expression is observed more centrally, in the underlying mesenchyme (B-C) (Cooper et al., 2018). Whole mount immunofluorescence revealed medial local expression of SHH associated with proliferative denticle placodes (D). Chicken feather buds display comparable patterns of gene expression at E10 (E-H), with epithelial expression of *Shh* in the leading tip of the feather bud (E), and dermal expression of both *fgf4* and *bmp4* (F-G) (Jung et al., 1998). Anti-SHH is also observed localised to individual developing feather buds (H).

As previously noted from section *IS*H (Fig. 3A), whole mount *IS*H reveals that expression of *shh* is restricted to the posterior facing apex of denticles undergoing morphogenesis (Fig. 7A). Conversely, expression of both *fgf4* and *bmp4* appears restricted to the mesenchymal papilla of the developing denticle, located behind the apical *shh* signal (Fig. 3B-C) (Cooper et al., 2018). Immunofluorescence reveals medial localisation of SHH in the denticle placode, within a highly proliferative epithelium (Fig. 7D). Comparable expression patterns are also observed in the chicken embryo, with feather buds expressing apically concentrated *Shh*, and dermal expression of both *Fgf4* and *Bmp4* within the feather bud papilla (Fig. 7E-G) (Jung et al., 1998). Here, immunofluorescence reveals an accumulation of SHH at the feather bud edge (Fig. 7H), potentially contributing to polarised outgrowth. Overall, we observe notable similarity of gene expression patterns during the early development of both shark denticles and chicken feathers, indicating that the conserved putative oGRN proposed here (Martin et al., 2016) is likely present across the skin appendages of phylogenetically distinct vertebrates.

## Discussion

This study has revealed several findings regarding the development of shark dermal denticles. First, during denticle emergence, many genes associated with tooth development are expressed in domains comparable to developing teeth, indicating that a conserved oGRN regulates the development of both of these skin appendages (Martin et al., 2016). This is particularly apparent in the epithelial tip of the denticle, which, in common with the shark dentition, shows reduced cell proliferation accompanied by expression of *shh, fgf3* and *mk* (Fig. 3). This provides further evidence to support the ancestral conservation of a modular cusp-making signalling centre analogous to the mammalian enamel knot (Thiery et al., 2022). This study therefore reveals a significant degree of serial patterning conservation between denticles and teeth.

Furthermore, our comparison of spatial gene expression patterns between the oral teeth and dermal denticles of sharks has implications for current models of odontode evolution. With the expression of identical genes observed between both oral teeth, which develop within a dental lamina, and external skin denticles (Fig. 5), our findings support the ‘inside and out’ model of tooth origin, suggesting the simultaneous deployment of a shared and co-opted oGRN (Smith and Coates, 1998, Fraser et al., 2010, Donoghue and Rucklin, 2016, Martin et al., 2016). Although our study is not intended to provide a framework for elucidating the origin of teeth, these data expand our knowledge of a shared and equivalent odontode-related genetic toolkit.

A notable difference between dermal denticles and oral teeth in sharks is the differential expression of *sox2* (Fig. 5), restricted exclusively in the dentition, to the cells of the dental lamina that contribute to the early dental epithelium and dental stem cell niche (Martin et al., 2016). This discrepancy underlies the difference in regenerative potential between the oral and dermal epithelia, with oral teeth exhibiting a continuous replacement system, in contrast to dermal denticles, which show no corresponding evidence of autonomous renewal (Martin et al., 2016). Although our findings generally conform to both the ‘inside-out’ and ‘inside and out’ models, both of which propose odontogenic potential to be universal to the oral and dermal epithelia, this differential expression of *sox2* is indicative of a deviation in the shared oGRN (Donoghue and Rucklin, 2016, Smith and Coates, 1998).

Genes expressed during both tooth and denticle development are also observed in the development of elasmobranch sensory receptors. This highlights similarities between these skin appendages, again suggesting a potential common evolutionary origin. This conforms to the ‘inside and out’ model which postulates that teeth and denticles may be derived from the collaboration of a precursory epithelial unit, perhaps with a sensory function, and a newly acquitted neural crest-derived cell type, which together developed odontogenic potential (Fraser et al., 2010, Baker and Bronner-Fraser, 1997). Progressive remodulation of common signals may then have led to the appearance and subsequent diversification of teeth and denticles (Fraser et al., 2010). Furthermore, in concurrence with previous research (Di-Poï and Milinkovitch, 2016, Cooper et al., 2017, Harris et al., 2008), our comparison of denticle and feather development (Fig. 7) demonstrates that the conservation of placode initiation and early morphogenesis is observed not only between the shark sensory system and odontodes, but also across skin appendages of phylogenetically distinct vertebrates. Later divergence in developmental processes then gives rise to the plethora of diverse skin appendages observed in nature.

We have characterised and compared the molecular development of odontodes in the shark (Fig. 3, Fig. 4, Fig. 5). However, there is growing evidence to suggest that, in addition to genetic regulation, mechanical systems are also paramount in controlling skin appendage emergence (Shyer et al., 2017, Shyer et al., 2013). This includes processes such as mechanosensation resulting from cellular aggregation, which can initiate genetic signalling cascades (Ho et al., 2019). In fact, integrated molecular and mechanical systems are known to control the precise patterning of the feather array in avian species (Ho et al., 2019). To obtain a comprehensive understanding of the both the physical and molecular systems at play during the development of both shark denticles and teeth, such mechanical processes should also be investigated in future studies. This will provide evolutionary perspectives regarding the emergence of such mechanical systems in the context of skin appendage development.

## Conclusions

Here, we have charted the development of dermal denticles, comparing conserved gene expression patterns with those of the developing shark dentition. This has revealed shared developmental processes across both shark denticles and sensory receptors, likely common to an ancient odontode. Patterns of both cell proliferation and specific gene signalling during placode initiation and subsequent morphogenesis imply the presence of a modular signalling centre comparable to the mammalian enamel knot (EK) (Vaahtokari et al., 1996), and the apical epithelial knot (AEK) in sharks (Thiery et al., 2022). Integration of these gene expression patterns with additional markers from tooth development add to the wider putative ancestral oGRN model (Fig. 4E) (Martin et al., 2016, Rasch et al., 2016). This generally conforms to the ‘inside and out’ model of tooth origin, which views the development of teeth and denticles as conserved at the molecular level (Smith and Coates, 1998, Fraser et al., 2010). However, the differential expression of the dental stem cell marker *sox2*, restricted to the epithelia of the oral dentition, implies a key difference in odontode initiation and regenerative potential between oral and epidermal locations. Future studies including RNA sequencing will yield a more comprehensive understanding of the molecular development of both dermal denticles and oral teeth, and elucidate how differential gene expression patterns are related to variation not only in their regenerative capacities, but also variation in their supporting tissues and attachment mechanisms.

## Acknowledgements

We thank Alexandre Thiery, and past and present members of the Fraser Lab for stimulating discussion associated with this project. We also extend our gratitude to Kyle Martin, Zerina Johanson (Department of Earth Sciences, Natural History Museum, London), Farah Ahmed, and Armin Garbout (Imaging and Analysis Centre, Natural History Museum, London) for assistance with micro-CT imaging. This research was supported by the following research grants: National Science Foundation IntBIO Collaborative Research grant: 2128032 (to G.J.F); Natural Environment Research Council (NERC) Standard Grant NE/K014595/1 (to G.J.F.); NERC PhD studentship (to L.J.R.), and Leverhulme Trust Research Grant RPG-211 (to G.J.F.). This work was also funded through “Adapting to the Challenges of a Changing Environment” (ACCE), a NERC-funded doctoral training partnership (to R.L.C.) ACCE DTP (NE/L002450/1). The authors declare that they have no conflict of interest.

